# GAMIBHEAR: whole-genome haplotype reconstruction from Genome Architecture Mapping data

**DOI:** 10.1101/2020.01.30.927061

**Authors:** Julia Markowski, Rieke Kempfer, Alexander Kukalev, Ibai Irastorza-Azcarate, Gesa Loof, Birte Kehr, Ana Pombo, Sven Rahmann, Roland F Schwarz

## Abstract

**Motivation:** Genome Architecture Mapping (GAM) was recently introduced as a digestion- and ligation-free method to detect chromatin conformation. Orthogonal to existing approaches based on chromatin conformation capture (3C), GAM’s ability to capture both inter- and intra-chromosomal contacts from low amounts of input data makes it particularly well suited for allele-specific analyses in a clinical setting. Allele-specific analyses are powerful tools to investigate the effects of genetic variants on many cellular phenotypes including chromatin conformation, but require the haplotypes of the individuals under study to be known a-priori. So far however, no algorithm exists for haplotype reconstruction and phasing of genetic variants from GAM data, hindering the allele-specific analysis of chromatin contact points in non-model organisms or individuals with unknown haplotypes.

**Results:** We present GAMIBHEAR, a tool for accurate haplotype reconstruction from GAM data. GAMIBHEAR aggregates allelic co-observation frequencies from GAM data and employs a GAM-specific probabilistic model of haplotype capture to optimise phasing accuracy. Using a hybrid mouse embryonic stem cell line with known haplotype structure as a benchmark dataset, we assess correctness and completeness of the reconstructed haplotypes, and demonstrate the power of GAMIBHEAR to infer accurate genome-wide haplotypes from GAM data.

**Availability:** GAMIBHEAR is available as an R package under the open source GPL-2 license at https://bitbucket.org/schwarzlab/gamibhear

Maintainer

## 1 Introduction

Genome Architecture Mapping (GAM) is a novel digestion- and ligation-free experimental technique for assessing the 3D chromatin structure from sequencing a collection of thin cryosectioned nuclear profiles (NuPs) (Beagrie *et al.*, 2017). Chromatin contacts between DNA loci can be inferred from the frequency at which loci are captured in the same NuP. One advantage of GAM over competing methods, such as Hi-C (Lieberman-Aiden *et al.*, 2009), is that GAM only requires several hundreds of cells to obtain high-resolution contact maps (Kempfer and Pombo, 2019; Beagrie *et al.*, 2020; Fiorillo *et al.*, 2020). This makes GAM particularly useful for the study of chromatin contacts in rare biological materials, such as human biopsies. Recently, there has been increasing interest in the allelespecific analysis of chromatin contacts, for which haplotyping, i.e. phasing of single nucleotide variants (SNVs) is key (Rivera-Mulia *et al.*, 2018; Cavalli *et al.*, 2019), but so far no algorithm exists for haplotype reconstruction from GAM data.

De-novo phasing is traditionally achieved through read-based methods such as HapCut, WhatsHap or HapCHAT (Bansal and Bafna, 2008; Patterson *et al.*, 2015; Edge *et al.*, 2017; Beretta *et al.*, 2018). In these methods, variants of the Minimum Error Correction (MEC) problem are used with varying error distributions and insert lengths (Bansal and Bafna, 2008). MEC views the given data (a fragments by SNV sites matrix of observed allele states) as potentially erroneous and asks for the least invasive way to correct the observations to enable conflict-free phasing. The MEC problem is computationally hard under a variety of conditions (Bafna *et al.*, 2005; Cilibrasi *et al.*, 2005). As a heuristic, HapCut converts MEC to a maximum cut problem and originally allowed for only single base pair errors (Bansal and Bafna, 2008). Selvaraj et al. (2013) later leveraged chromosome territories (Meaburn and Misteli, 2007) and extended HapCut to Hi-C data by accommodating Hi-C specific h-trans errors. H-trans errors are haplotype switch errors that occur when a genomic region interacts with another genomic region located on the other homologous chromosomal copy (in trans). HapCut2 now includes population-based statistical phasing (Bansal, 2019) and implements a variety of different error models to accommodate different sequencing technologies (Edge *et al.*, 2017).

Alternative formulations to the phasing problem seek to partition the observed fragments (Duitama *et al.*, 2010), or the aggregated co-occurrence frequencies of SNVs (Tourdot and Zhang, 2019), into two classes corresponding to the two haplotypes by minimising a measure of inconsistency. To facilitate haplotype reconstruction from GAM data, we here also use an aggregation step and formulate the problem on co-occurrence evidence derived from the raw GAM NuPs. This formulation is equivalent to finding a ground state of a spin glass system (in statistical physics), which is equivalent to a maximum cut problem (Tourdot and Zhang, 2019).

The resulting algorithm GAMIBHEAR (GAM-Incidence Based Haplotype Estimation And Reconstruction) employs a graph representation of the co-occurence of SNV alleles in NuPs for whole-genome phasing of genetic variants from GAM data. GAMIBHEAR accounts for the GAM-specific probabilities in capturing parental chromosomal segments as part of the random cryosectioning process. We assess the performance of GAMIBHEAR on the hybrid mouse embryonic stem cell line F123 with known haplotype structure. Despite the sparsity of GAM data, GAMIBHEAR allows for accurate, dense, genome-wide haplotype reconstruction.

## 2 Methods

### 2.1 Definitions, problem statement and objective

Our goal is to reconstruct haplotypes from GAM data. A sequenced GAM dataset consists of reads from many nuclear profiles (NuPs). Each NuP is the result of random sectioning of the nucleus and captures ultra-sparse *local* sequence information, where *local* refers to genomic loci in close proximity in the 3D arrangement of the genome, including but not limited to loci proximal in linear distance. Thus, reads from single NuPs cover a small proportion of the whole genome with consecutive stretches of genomic DNA that reflect chromatin looping in and out of a thin nuclear slice (illustrated in Figure 2B). Our main assumption here is that alleles of any two heterozygous SNVs captured in a nuclear slice are likely to originate from the same parental copy, and that this likelihood decreases with increasing genomic distance between the two SNVs.

**Figure 1:**
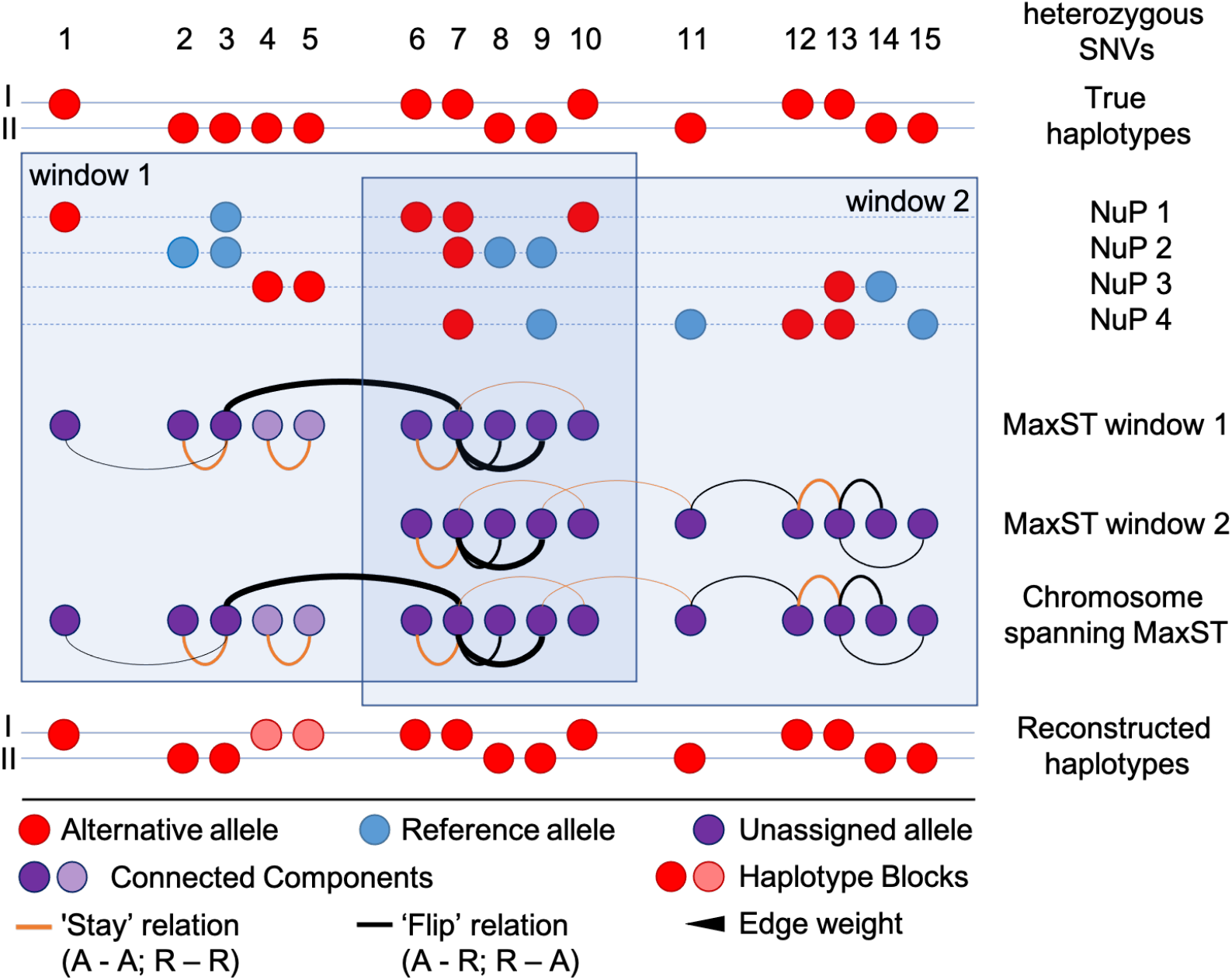
Schematic overview of the graph phasing algorithm. The location of alternative alleles of heterozygous SNVs on the two parental chromosomes describes the true haplotypes (top). NuPs 1 - 4 are sparse local samples of the true haplotype structure. At heterozygous SNV positions either the alternative (red) or reference allele (blue) can be observed. In overlapping windows, graphs of co-observed SNVs are built over all NuPs. Edges are of either stay (orange) or flip (black) type and edge weights correspond to the co-observation frequency (line width) and are optionally proximity-scaled. A set of SNVs that is itself not co-observed with other SNVs in the same window forms its own connected component in the graph (e.g. SNV 4 and SNV 5 in NuP 3, window 1). MaxSTs (forests in case of multiple connected components) are calculated per window and combined to yield a sparse but chromosome-spanning graph. The MaxST of this sparse graph is used to assign alternative alleles to the final reconstructed haplotypes (bottom). Connected components in the final MaxST form separate, possibly nested haplotype blocks (red/pink). As the leftmost SNV of each separate haplotype block is assigned to haplotype 1, SNVs 4 and 5 are correctly phased relative to each other (stay relation), but assigned to the wrong haplotype.

**Figure 2:**
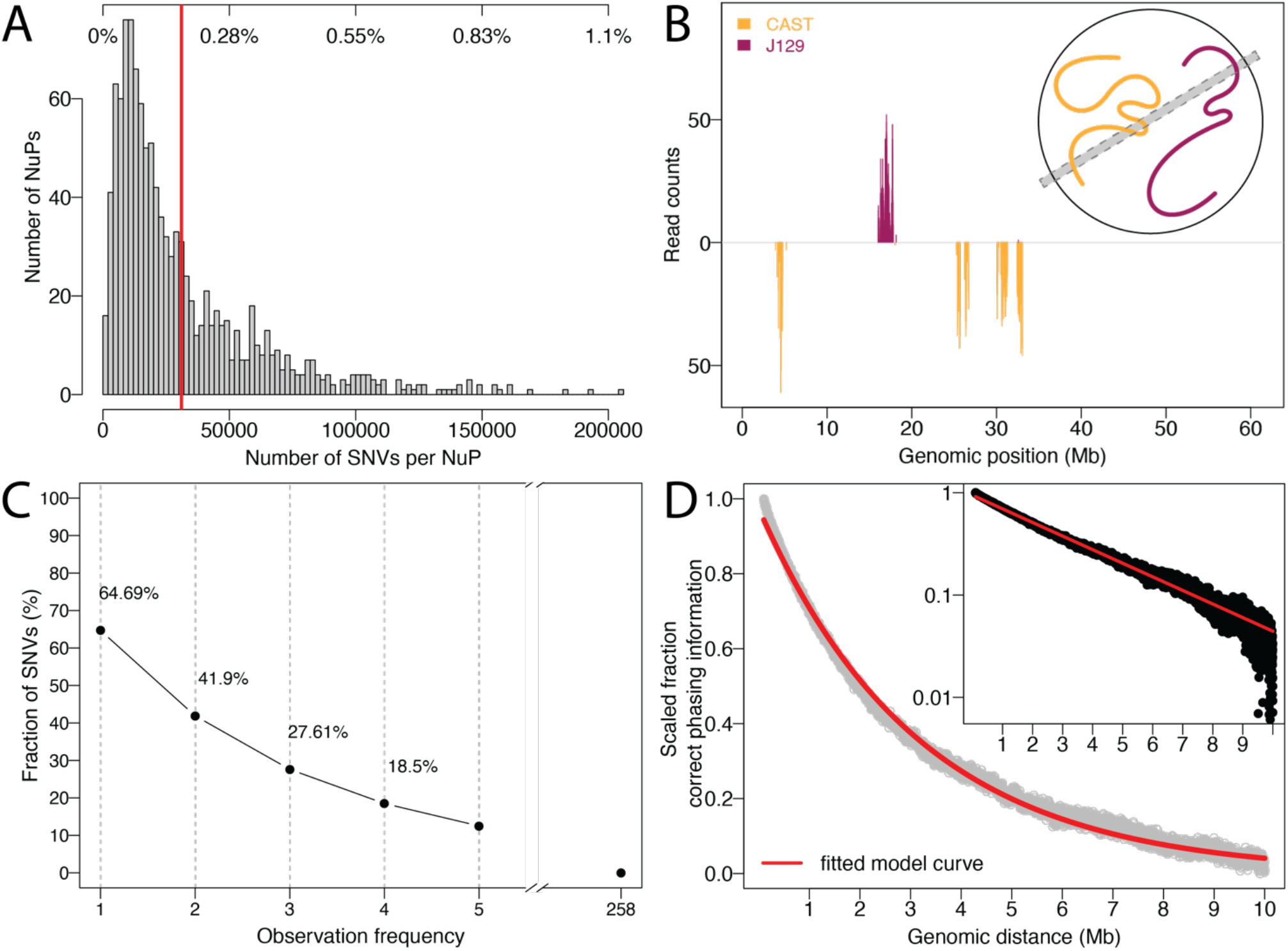
GAM captures local phasing information: **A)** Histogram of the number of observed SNVs per NuP in the F123 dataset (fraction of all SNVs at top, mean = 0.171%, red line). **B)** Example of read counts supporting the CAST (orange, downwards) and J129 (red, upwards) alleles in a single NuP on chromosome 19, visualising the sparsity of GAM data. Inset depicts physical capturing of respective genomic regions in a slice (grey area) by cryosectioning in a GAM experiment. **C)** Cumulative fraction of SNV observation frequencies. 64.69% of SNVs are observed at least once, 41.9% of SNVs are observed at least twice across all NuPs. **D)** The fraction of correct phasing information decreases exponentially with increasing genomic distance of observed SNV pairs. The fit of the exponential curve to the fraction of correct phasing information of SNV pairs with genomic distance between 1 bp and 10 Mb is shown in red. The inset shows the decrease of correct phasing information on a logarithmic scale.

We assume that the set of heterozygous SNVs is given and that the SNV alleles have been determined per NuP. Let *N* be the number of NuPs and *M* be the number of heterozygous SNVs in the genomic region of interest (e.g., a chromosome or chromosome arm; sites with homozygous SNVs are ignored). Then the problem input is a ternary *N* × *M* matrix *D* with *D_ij_* = 1 if the reference allele is observed in NuP *i* at SNV site *j, D_ij_* = −1if the alternative allele is observed, and *D_ij_* = 0if there is no unique observation (e.g. due to lack of coverage or if both alleles are observed in the same NuP).

The goal is to reconstruct the two haplotypes (allele states on the same parental copy). Formally, a haplotype is a vector *h* ∈ {−1, 1}^*M*^ with *h_j_* = 1 if the reference allele is found at site *j* and *h_j_* = −1 for the alternative allele. One of the two haplotypes *h* determines the other one as –*h*.

The GAM input data in principle contains the information to infer*h*. Consider the relation between SNV sites *j* and *k* in NuP *i*. The two sites can be in a “flip” relation, where the alternative (alt) allele (−1) of one site is observed with the reference allele (+1) of the other site (product*D_ij_* · *D_ik_* = −1), and a “stay” relation, where both SNVs show either the reference or alternative allele (product *D_ij_* · *D_ik_* = 1).

We thus compute the *M* × *M evidence matrix A*:= *D^T^D*, which contains the accumulated counts of the stay-flip relations summed over all NuPs, i.e., 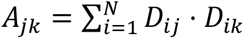, such that positive values indicate more stay observations (*A_jk_* > 0: ‘stay’ between sites *j* and *k*; *j,k* = 1,.…, *M*) and negative values indicate more flip observations (*A_jk_* < 0: ‘flip’ between sites *j* and *k*). An equal number of observed stays and flips leads to zero entries (*A_jk_* = 0).

The goal of the haplotype reconstruction algorithms we develop here is to find *h* using information contained in *A*: If *A_jk_* > 0, then we should have *h_j_* = *h_k_*, and if *A_jk_* < 0, then *h_j_* = −*h_k_*. However, the information in *A* may be conflicting when considering transitivity: Consider three sites *j, k, l* with *A_jk_* > 0, *A_jl_* > 0, *A_jl_* < 0. Thus, decisions need to be made on how to resolve conflicting information in the evidence matrix *A*.

We formulate the problem as follows: given the *M × M* matrix *A*, we seek *h* ∈ {−1, 1}^*M*^ to

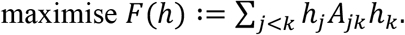

This formulation encourages *h_j_* and *h_k_* to take the same sign if *A_jk_* > 0 and different signs if *A_jk_* < 0. This maximization problem is equivalent to finding an exact ground state for a spin glass in physics and is known to be NP-hard in general and can be cast as a maximum cut problem on a graph induced by *A* (Tourdot and Zhang, 2019). Here we propose heuristic algorithms that make use of known properties of the evidence matrix *A* (potentially proximity-scaled; see below) and evaluate them against a dataset with a known correct solution.

Before we state two such algorithms, let us first relax our notion of what we accept as a solution. Above, we defined a (fully resolved) haplotype as a vector *h* ∈ {−1, 1}^*M*^with *h_j_* = 1 if the reference allele is found at site *j* and *h_j_* = −1for the alternative allele. However, the available data may not be sufficient to fully resolve the haplotype. Where no phasing information is available, we allow partial solutions (“blocks”) as follows. Let *J*:= (*J*_1_, *J*_2_,…, *J_K_*) be a partition (disjoint union) of {1,…, *M*}into *K* blocks. Then a solution of the GAM haplotype reconstruction problem for input matrix *D* with partition *J* is a collection of *K* binary vectors *h*^1^ ∈ {−1,1}^*J*_1_^,…, *h^K^* ∈ {−1, 1}^*J_K_*^. Each of the *K* blocks is solved independently, and no statement is made about the connection between these blocks. The blocks are often intervals, but may be arbitrary subsets of all sites, especially for GAM data. Obviously, solutions with fewer independent blocks are more desirable.

### 2.2 Haplotype reconstruction algorithms

#### 2.2.1 Neighbour phasing

We first consider a baseline phasing strategy that leverages the most reliable short-range haplotype information on neighbouring SNVs only (“neighbour phasing”). In the above notation, we only consider the first off-diagonal of *A*, i.e., *A*_*j*,(*j*+1)_ for *j* = 1,…, *M*. Essentially, this resolves possible conflicting information by ignoring a large fraction of the available data, and only considering a single path between any two sites *j* ≤ *k*: *j* → *j* + 1 → ··· → *k*. The reconstructed haplotype starts (arbitrarily) with the reference allele, thus *h*_1_ = 1. Once *h_j_* is determined, we set *h*_*j*+1_: = *h_j_*·sign(*A*_*j*,(*j*+1)_), i.e. we stay or flip according to the sign of *A*_*j*,(*j*+1)_. In case of a tie or when SNV*j* and *j*+1 are never co-observed in the same NuP (*A*_*j*,(*j*+1)_ = 0), we start a new independent haplotype block where *h*_*j*+1_ = 1. Solutions produced by neighbour phasing consist of blocks that are intervals. The resolved blocks can be expected to be correct with high probability, but also short, and therefore of limited use.

#### 2.2.2 Graph phasing with optional proximity scaling

We extend the local proximity of SNVs from immediate neighbours to larger genomic windows using a graph-based approach (Figure 1). To improve computational efficiency each chromosome is divided into windows of a fixed number *L* of SNV sites with half a window size overlap. Phasing is carried out on each window independently and results per window are subsequently reconciled (see below). To process a window, we restrict the *N* × *M* input matrix *D* = (*D_ij_*) to the window’s sites and only consider the reduced *N* × *L* matrix *D* and the derived *L* × *L* evidence matrix *A* = (*A_jk_*). We systematically evaluated different windows sizes in terms of runtime, memory usage and phasing completeness and accuracy. We settled on *L* = 20,000 SNVs as the default, as it causes only a marginal reduction in accuracy while improving completeness and drastically reducing computational demands (see Supplementary Note S6).

As we assume that the reliability of phasing information within a NuP decreases with genomic distance, we include an option to scale the information in *A* element-wise by a weight matrix *W* = (*W_jk_*), where *W_jk_* depends on the genomic distance *d_jk_* between sites *j* and *k*. We use a simple exponential decay model, where *W_jk_* = *C*·exp (−λ *d_jk_*) for *d_jk_* in a certain range [*D_min_, D_max_*], and *W_jk_* = 1 for *d_jk_* < *D_min_* and *W_jk_* = 0 for *d_jk_* > *D_max_*. The choice of appropriate parameters *C* > 0, λ > 0 and 0 ≤ *D_min_* < *D_max_* is discussed below. In the following, *A* represents the *proximity-scaled evidence matrix* (*A_jk_* ← *W_jk_* · *A_jk_*).

At this point, there are four potential reasons for *A_jk_* = 0: First, sites *j* and *k* may never co-occur in any NuP. Second, they may never be considered in the same window of *L* sites. Third, their genomic distance may be larger than *D_max_*. Fourth, an equal number of observations of stay and flip relations may be encountered between sites *j*and *k*.

The non-zero entries in *A* induce an undirected weighted graph. Its *L* vertices are the sites of the current window. An edge between sites *j* and *k* exists with weight *A_jk_* if *A_jk_* ≠ 0. Two sites in the same connected component of this graph are typically connected by many paths. Consider a single arbitrary path between sites *j* and *k*. The number of negative-weighted edges along the path determines the haplotype assignment: if the number is even, then *h_k_* = *h_j_*; if it is odd, then*h_k_* = −*h_j_*. Different paths between the two sites can be conflicting in their haplotype assignment. However, if the graph is reduced to a tree (or forest in case of more than one connected component), there is a unique path between each pair of sites (in the same connected component). Because the absolute values |*A_jk_*| indicate strength of direct evidence for the relation between sites *j*and *k*, we compute a *maximum spanning tree* (MaxST) of each connected component based on absolute edge weights |*A_jk_*| using Kruskal’s algorithm. Recall that the problem is solved on (potentially dense graphs of) windows, so the required running time is *O*(*L*^2^ log *L*)for each window. The MaxST has the property that the resulting path between any two sites *j* and *k* maximises the minimum weight of the path’s edges among all possible paths between *j* and *k* (Hu, 1961), so we construct the graph by maximising the weakest evidence link between each pair of sites of the window, which appears to be a reasonable heuristic for the given problem. The computed MaxST then determines the haplotypes (or set of haplotype blocks in case of a forest of MaxSTs) for the current window.

To infer haplotypes across the whole chromosome, the MaxSTs of overlapping windows must then be joined into a chromosome-wide graph. For this, we join the (overlapping) MaxSTs of all windows into a new graph consisting of all *M* SNV sites as nodes and the union of edges of all MaxSTs. Because each node is in at most two MaxSTs, the number of edges in the union is bounded by 2(*M* – 1). In order to solve possible disagreements stemming from the results of overlapping windows in this sparse graph, we again determine a MaxST (if necessary, on each connected component separately) in *O*(*M log M*) time to obtain a unique path between any two connected sites.

For the output, each connected component defines an independent haplotype block. The haplotype of the leftmost SNV site *h*_1_ (with smallest genomic coordinate) in each block is arbitrarily set to *h*_1_: = 1, and the other states *h_j_* are computed according to the number of negative-weighted edges on the unique MaxST path between the first site and *j*.

Including phasing information from non-adjacent SNV pairs will improve completeness and yield larger, potentially chromosome-spanning haplotype blocks. In the reconstructed haplotypes of the graph phasing approach, blocks can be nested. The inclusion of phasing information from more distant SNV pairs might compromise the overall accuracy of the results, however the proximity scaling is expected to keep the introduction of misleading information to a minimum.

### 2.3 Performance measures

To assess the quality of the reconstructed haplotypes we compare GAMIBHEAR estimates with the haplotypes of the F123 mouse embryonic stem cell (mESC) line obtained from whole-genome sequencing of the parental mouse strains (see Supplementary Note S1). The overall quality of reconstructed haplotypes depends on both the completeness of the reconstructed haplotype blocks as well as the phasing accuracy of the SNVs contained.

In terms of completeness we report the total proportion of phased heterozygous SNVs next to the standard S50 (Lo *et al.*, 2011), N50 (Lander *et al.*, 2001) and adjusted N50 (AN50; (Lo *et al.*, 2011)) metrics which give an impression of the median size (in SNVs) and span (in bp) of the reconstructed haplotype blocks. To enable comparisons with previous phasing approaches of the F123 cell line (Selvaraj *et al.*, 2013) we report the metrics in percent of the phasable variants (number of input variants) and phasable genome (range between leftmost SNV and rightmost SNV per chromosome, 97,58% of the genome), respectively.

To evaluate accuracy, we report the Switch Error Rate (SER), defined as the proportion of adjacent variant pairs that were phased incorrectly out of all phased variant pairs. We also report the adjusted Switch Error Rate (adjusted SER) to account for incomplete or fragmented phasing results, by penalising unresolved transitions between adjacent variant pairs with 0.5 switch errors, to account for, on average, a 50% chance of assigning the wrong phase. Fragmented phasing occurs when the phasing graph is composed of many small components with phasing information within but not between components. Additionally, the global haplotype agreement is reported, calculated by direct comparison of the reconstructed and true haplotypes (i.e. alt-ref configurations). To be able to relate the results to the size of the GAM dataset, we also report the quality of haplotypes reconstructed from incrementally increasing subsets of 100 NuPs chosen at random in ten iterations (Figure 3). All performance measures are given in averages across all 19 chromosomes. For a more detailed description and motivation of the individual metrics please see Supplementary Note S4.

**Figure 3:**
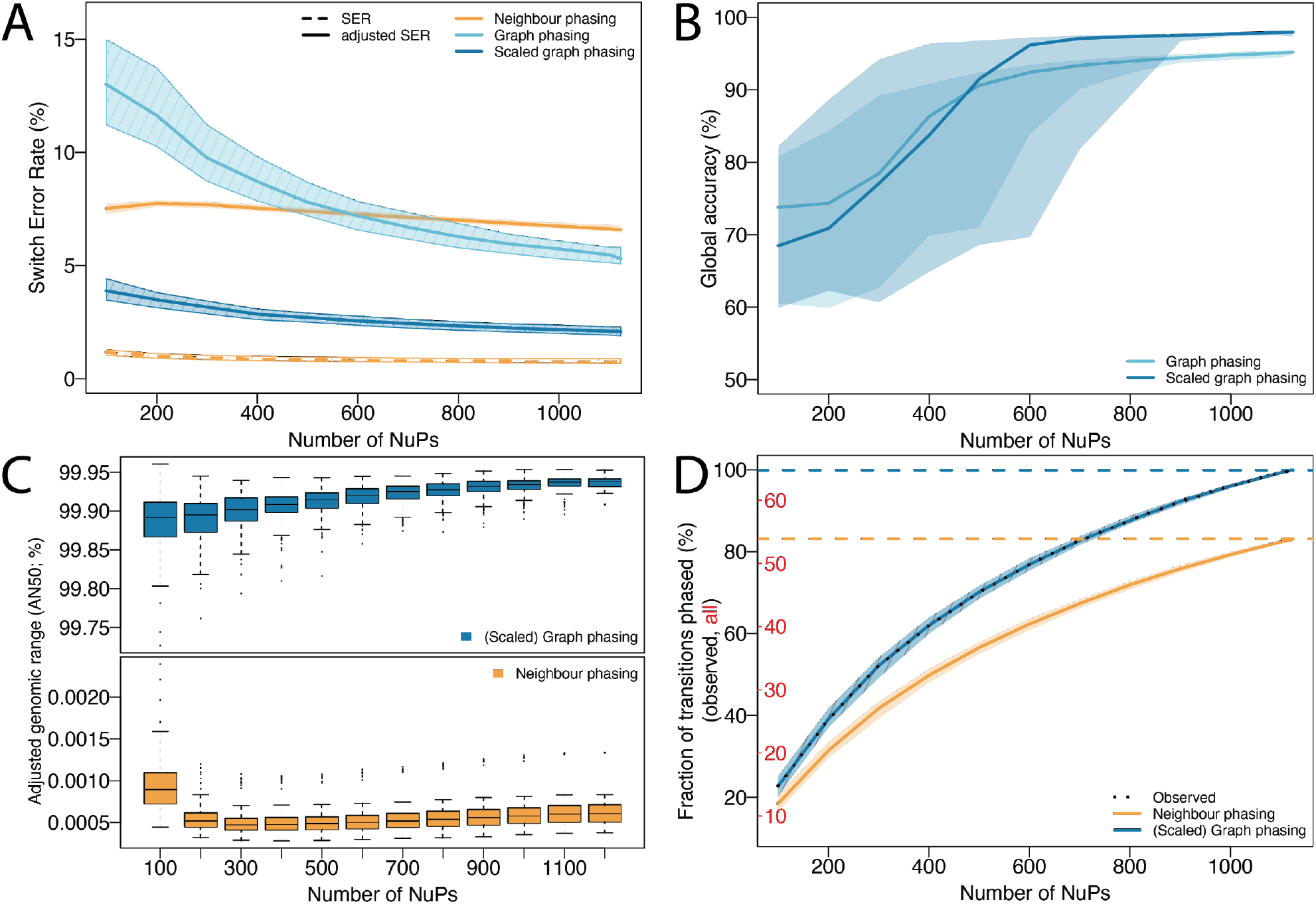
**Quality of reconstructed haplotypes** for neighbour phasing (orange), basic graph phasing (light blue) and proximity-scaled graph phasing (dark blue) for an increasing number of NuPs. Lines show the median value, shaded areas indicate the interquartile range of results across all chromosomes. **A) Local accuracy (SER)**: In graph phasing, SER decreases with an increasing number of NuPs as more information becomes available. Neighbour phasing in contrast shows a low SER independent of sample size (dashed orange line) due to a small number of phased transitions which are accurate. Adjusted SER penalises unphased transitions and shows this difference: neighbour phasing performance (solid orange line) is substantially lower, graph phasing performance is unchanged (SER and adjusted SER lines overlap). Proximity-scaled graph phasing shows lowest adjusted SER overall. **B) Global Accuracy (haplotype agreement)** improves with increasing sample size and proximity scaling further improves performance. **C) Completeness (AN50):** Graph phasing reconstructs dense, nested chromosome-spanning blocks even for low sample sizes (top), independent of proximity scaling. Neighbour phasing yields a large amount of small unconnected adjacent blocks, which are never nested, thus N50=AN50 (bottom). **D) Completeness (% transitions phased)**: Percentage of transitions phased relative to all known SNVs (red) and all SNVs observed at least once in the full dataset (black, see Figure 2C). The number of observed SNVs and thus phasable transitions increases with increasing number of NuPs (dashed black line). Graph phasing predicts 99.96%, neighbour phasing predicts 83.02% of observed transitions.

### 2.4 GAMIBHEAR implementation

The presented haplotype reconstruction algorithms are implemented in the R package GAMIBHEAR. GAMIBHEAR is open source and freely available under the GPL-2 license at https://bitbucket.org/schwarzlab/gamibhear.

## 3 Results

### 3.1 Benchmark dataset

The F123 mouse embryonic stem cell line was derived from a hybrid F1 mouse resulting from the cross of the two inbred, homozygous mouse strain*s CAST (Mus musculus castaneus)* and *J129 (Mus musculus domesticus J129)*. The F1 generation is thus heterozygous at all loci for which their parents have different alleles. As the parental mouse strains are both fully sequenced, the haplotypes of the F123 cell line were derived from SNV sets called on the parental strains (see Supplementary Note S1). Its known haplotype makes the F123 cell line an ideal model for benchmarking phasing algorithms.

Using the GAM method 1281 single NuPs were generated from the F123 mESC cell line (available at 4D Nucleome Consortium data portal accession number 4DNBSTO156AZ), out of which 1123 passed quality control (unique 4DN identifiers provided in Supplementary Data). We extracted on average 305,377 reads from 1123 NuPs, covering 0.171% (± 0.167) of the 18,150,228 heterozygous SNVs per nuclear slice (Figure 2A); exemplary data of genomic regions captured in a single NuP is shown in Figure 2B. Out of all F123 SNVs, 11,741,055 (64.69%) were observed at least once, 7,605,321 SNVs (41.9%) were observed at least twice (Figure 2C). For more details on F123, the generation of the benchmark haplotypes and the data preprocessing see Supplementary Notes S1 – S3.

### 3.2 Exponential proximity scaling

Our method includes the option of exponentially downweighting evidence information *A_jk_* with increasing genomic distance (see Section 2.2.2). To validate this assumption and to choose optimal decay parameters, we examined the empirical probability *p* of two alleles coming from the same haplotype in the F123 data based on their genomic distance *d* and fit an exponential function *p* = *C* · *e*^−λ·(*d−D*_min_)^ using non-linear least squares. For this model we only considered pairs of sites within the interval [*D_min_, D_max_*]= [1 bp, 10 Mb], where the decay in phasing information is most pronounced (Figure 2D). The distance can be individually assigned by the user and probabilities 1 and 0 are assumed below *D_min_* and above *D_max_* respectively. Parameter *C* = 1 then describes the co-observation probability at a genomic distance of 1 bp with an exponential decay parameter of *λ* = 3.173 o 10^-7^. This simple exponential model describes the empirical distribution (Figure 2D) well and thus appears to be a good model for the reliability of the raw evidence as a function of genomic distance. In the following, we evaluate our graph phasing approach with and without proximity scaling.

### 3.3 Performance of GAMIBHEAR

#### 3.3.1 High quality haplotype reconstruction from GAM data

We evaluated the quality of the haplotypes reconstructed with GAMIBHEAR in terms of completeness and accuracy by comparing results to the true haplotypes of the F123 cell line (Table 1).

**Table 1:**
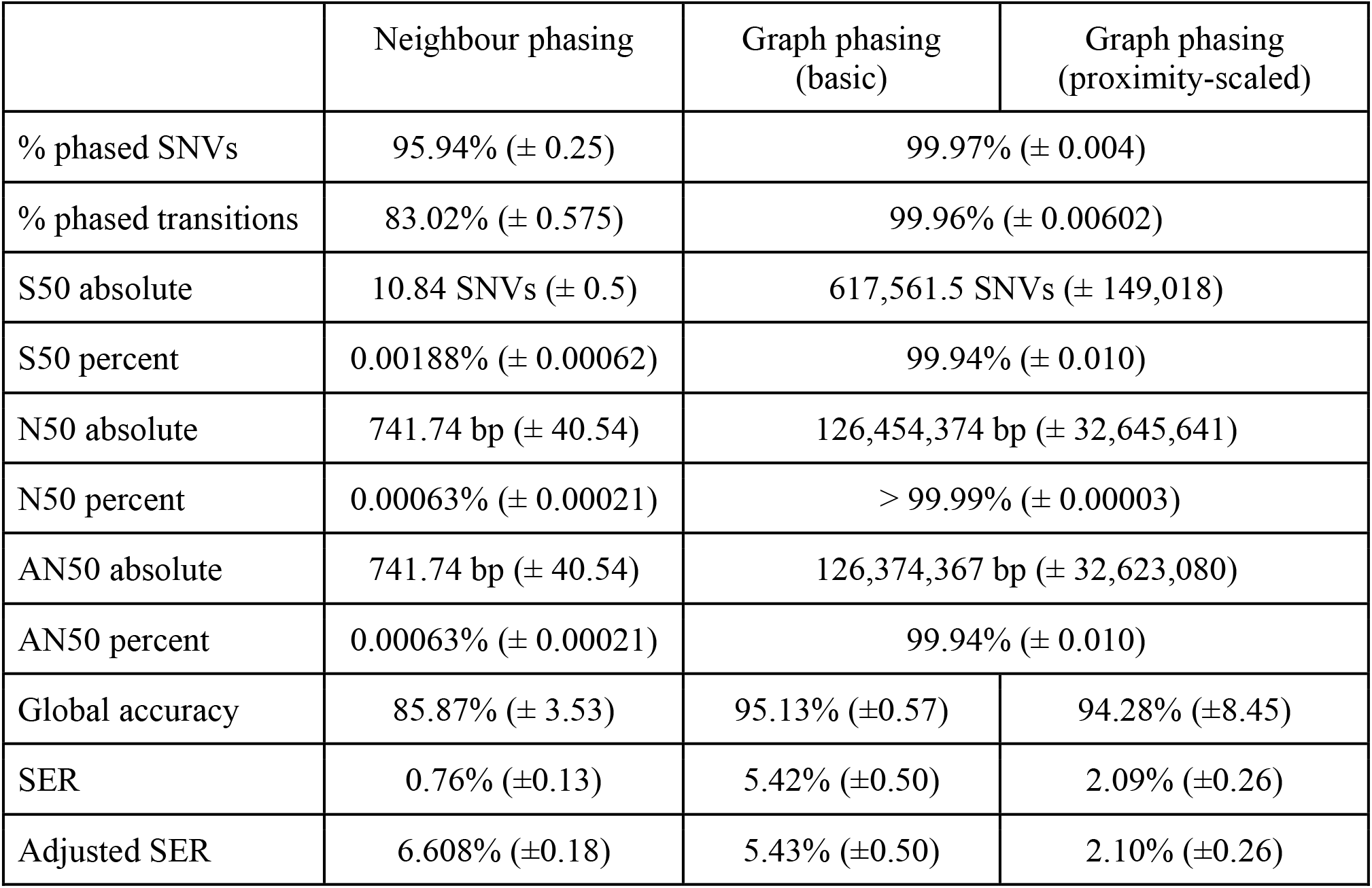
**Comparison of quality measures** for the neighbour phasing algorithm, basic and proximity-scaled graph phasing algorithm for the full dataset. The mean of per-chromosome values is reported, standard deviation in brackets. Percent phased SNVs and transitions are reported in relation to observed SNVs. For a per chromosome report of accuracy results see Supplementary Note S5.

|  | Neighbour phasing | Graph phasing<br>(basic) | Graph phasing<br>(proximity-scaled) |
| --- | --- | --- | --- |
| % phased SNVs | 95.94% ($\pm 0.25$) | 99.97% ($\pm 0.004$) | |
| % phased transitions | 83.02% ($\pm 0.575$) | 99.96% ($\pm 0.00602$) | |
| S50 absolute | 10.84 SNVs ($\pm 0.5$) | 617,561.5 SNVs ($\pm 149,018$) | |
| S50 percent | 0.00188% ($\pm 0.00062$) | 99.94% ($\pm 0.010$) | |
| N50 absolute | 741.74 bp ($\pm 40.54$) | 126,454,374 bp ($\pm 32,645,641$) | |
| N50 percent | 0.00063% ($\pm 0.00021$) | > 99.99% ($\pm 0.00003$) | |
| AN50 absolute | 741.74 bp ($\pm 40.54$) | 126,374,367 bp ($\pm 32,623,080$) | |
| AN50 percent | 0.00063% ($\pm 0.00021$) | 99.94% ($\pm 0.010$) | |
| Global accuracy | 85.87% ($\pm 3.53$) | 95.13% ($\pm 0.57$) | 94.28% ($\pm 8.45$) |
| SER | 0.76% ($\pm 0.13$) | 5.42% ($\pm 0.50$) | 2.09% ($\pm 0.26$) |
| Adjusted SER | 6.608% ($\pm 0.18$) | 5.43% ($\pm 0.50$) | 2.10% ($\pm 0.26$) |

##### Neighbour phasing performance

The neighbour phasing algorithm was built to exploit the most reliable short-range haplotype information of neighbouring co-observed SNVs, at the expense of completeness. This conservative algorithm shows the lowest switch error rate (SER) of the reconstructed haplotypes (0.76%, Figure 3A), demonstrating strong local phasing information in GAM data. However, although over 95% of input SNVs were phased into adjacent haplotype blocks of at least size 2, the number of independent blocks is high (on average 79,965 blocks per chromosome), their size is small (Figure 3C) and thus only 83% of possible transitions between neighbouring SNVs could be phased (Figure 3D). Median haplotype blocks connect less than 11 SNVs (S50, 0.00188% of the phasable SNVs) and span less than 742 bp (N50, 0.00063% of the phasable chromosome), showing drastically low completeness. This low completeness is evident in the stark contrast between SER (0.76%) and adjusted SER (6.61%), confirming that neighbour-phasing yields small locally constrained but accurate phasing blocks (Figure 3A). These locally accurate haplotypes confirm the presence of a strong local phasing signal in GAM data, but do not yield accurate phasing genome-wide. This algorithm shows the lowest global haplotype accuracy of 85.87%.

##### Graph phasing performance

The additional higher-order phasing information considered by the graph phasing algorithm substantially improves the completeness of the reconstructed haplotypes independent of proximity scaling (Figure 3C). Over 99.9% of input SNVs were phased into haplotype blocks, over 99.9% of them into one main haplotype block (S50), spanning more than 99.99% of the phasable genome (N50) and phasing 99.96% of transitions (Figure 3D). Adjusting the span of the largest block by the fraction of SNVs phased within yields an AN50 value of over 99.9% (Figure 3C). The graph phasing algorithm thus reconstructs dense chromosome-spanning haplotypes. Considering larger SNV windows increases the risk of integrating incorrect phasing information from co-observed SNV pairs located on homologous chromosome copies. Consequently, the accuracy of reconstructed haplotypes is lower compared to strict neighbour phasing. The basic graph phasing approach yielded results with ~5% SER (Figure 3A) and over 95% global accuracy (Figure 3B). To improve accuracy while maintaining completeness we introduced proximity scaling and successfully reduced SER to ~ 2% and increased global accuracy to ~ 98% (Figure 3 A and B) with the exception of a few outliers (see Supplementary Figure 2). Those outliers are caused by a single switch error occurring within a haplotype block, which inverts the assignment of subsequent alleles, formally reducing global accuracy while maintaining SER and high, reliable local accuracy. Since the graph phasing resulted in highly complete haplotypes with a very low number of haplotypes blocks (on average 76 blocks per chromosome), the SER adjusted for unphased transitions only showed negligible changes compared to the unadjusted SER (adjusted SER: unscaled: 5.43%, scaled: 2.10%). In conclusion, proximity-scaled graph phasing shows best performance overall and reconstructs accurate, chromosome-spanning haplotypes.

#### 3.3.2 Performance at lower SNV density

To show the effect of SNV density on the quality of haplotype reconstructions, we subsampled the F123 SNV set (~ 8 SNVs per 1kb) to resemble human SNV density (~ 1-1.5 SNVs per 1kb, (1000 Genomes Project Consortium *et al.*, 2015)) and evaluated the resulting haplotypes reconstructed using the best-performing proximity-scaled graph phasing algorithm (See Supplementary Note S7).

GAMIBHEAR reconstructed accurate, dense, chromosome-spanning haplotypes: 99.96% of input SNVs were phased, of which 99.95% are within the main, chromosome-spanning haplotype block. This block spans 100% of the phasable genome (97.56% of the full genome). The median global accuracy of 96.64% and the switch error rate of 4.84% show that the quality of the reconstructed haplotypes in a subsampled dataset is only slightly lower compared to the haplotypes reconstructed from the full dataset, indicating that the algorithmic approach is largely independent of SNV density and thus applicable to human data. GAMIBHEAR thereby showed greatly improved resolution at a slightly reduced global accuracy compared to HaploSeq on comparably downsampled data (~32% of input SNVs phased; 98.9% global accuracy) (Selvaraj et al. 2013) (Supplementary Note S9).

#### 3.3.3 Time and Memory usage

Phasing 11,741,055 heterozygous variants from the full 1123 NuP GAM dataset took approximately 1.5 hours and 16 GB (largest chromosome 1: ~ 14 min, 0.9 GB) using the neighbour phasing algorithm, ~ 5 h and 30 GB using the basic / proximity-scaled graph phasing algorithm (largest chromosome 1: ~ 27 min, 26 GB) with default settings on a desktop PC with 64 GB of RAM without parallelisation. However, computation can be carried out in parallel on multiple chromosomes for a further speed increase using the “cores” option. Reconstructing haplotypes from the dataset subsampled to human SNV density using the best performing proximity-scaled graph phasing algorithm took ~ 38 min and 20 GB for the whole genome (largest chromosome 1: ~ 3 min, 12 GB). For more details see Supplementary Note S6.

### 3.4 Comparison with existing methods

We compare GAMIBHEAR to the haplotype assembly methods WhatsHap (wMEC solver) (Patterson *et al.*, 2015) and HapCHAT (k-constrained MEC solver) (Beretta *et al.*, 2018), both designed for reconstructing haplotypes from long reads. For this we converted GAM NuPs into pseudo-long reads by adapting the ternary input matrix *D* (see section 2.2 and Supplementary Note S8). WhatsHap has a maximum coverage threshold of 23 reads which is exceeded in the F123 GAM data on a small number (0.0073%) of SNVs. This resulted in the read selection heuristic of WhatsHap to select only 69 of 1087 pseudo long reads (6.35%), thereby retaining only 11,039 SNVs (1.17% of input SNVs). In conclusion, coverage constraints in WhatsHap prevent its direct application to GAM data. Recently, HapCHAT was introduced to address this shortcoming by merging reads that are likely to originate from the same chromosome copy before read selection. In HapCHAT 1087 pseudo long reads were thus merged into 691 reads, 63 of which were selected for subsequent phasing, covering 604,358 SNVs (64.18% of input SNVs). From these, HapCHAT reconstructed a chromosome spanning haplotype block, with a global accuracy of 81.36% and an SER of 11.38% (compared to a global accuracy of 98.03% and SER of 1.98% using GAMIBHEAR). The MEC cost was reported as 307,734. This shows that in addition to the differences in coverage, the unique properties of GAM data prevent direct application of long read MEC solvers for phasing. For details see Supplementary Note S8.

## 4 Discussion

The phasing problem has been extensively studied and approaches to solve it are typically specific to and optimised for certain experimental designs and datatypes, such as Hi-C (Edge *et al.*, 2017) and long reads (Patterson *et al.*, 2015; Beretta *et al.*, 2018). Although both GAM and Hi-C capture the spatial proximity of SNVs in the nucleus, the coverage and error distributions of the GAM cryosectioning process are sufficiently different from those of Hi-C that existing MEC solvers are not directly applicable. In Hi-C, phasing information is contained in ligated chimeric reads of genomic loci harboring at least two SNVs, which can be very distant in linear genomic space but typically from the same chromosomal haplotype. In contrast, in GAM, phasing information is contained in NuPs, which yield individual short reads of both haplotypes and only maintain haplotype fidelity locally. Thus, in contrast to Hi-C, where h-trans errors remain rare, GAM NuPs frequently switch haplotypes. A Hi-C dataset furthermore consists of millions of reads, of which only a small percentage is useful for phasing as they rarely cover two SNVs or more (Giorgetti *et al.*, 2016). In contrast, a GAM experiment has in the order of 10^3^ NuPs, but a GAM NuP covers many SNVs (Figure 2B). A single NuP therefore contains many long stretches of haplotype-resolved SNVs that allow “neighbor phasing”, which is not available with Hi-C and which shows that phasing with Hi-C and GAM data are two distinct computational problems.

In addition, SNV coverage in GAM data varies greatly and non-uniformly, which interferes with MEC solvers for long read data that are fixed parameter tractable in the coverage and thus require the maximum coverage per SNV to be low (Patterson *et al.*, 2015). To ascertain these differences, we tested GAM data on the long-read MEC solvers WhatsHap and HapCHAT. HapCHAT only yielded SERs > 10%, owing to differences in the underlying technologies: long reads are not affected by haplotype switches but will frequently include single-nucleotide sequencing errors; GAM data, however, shows frequent switches in observed haplotypes, affecting all following SNVs. Due to these fundamentally different data characteristics MEC solvers designed for haplotype assembly from long reads yield unsatisfying results when employed on GAM data.

We did not attempt to transform GAM data for use with HapCut2, as it has been well known and stated by the authors that the performance of HapCut2 strongly depends on the correct error model being used and no such model exists for GAM data (Edge *et al.*, 2017).

The closest comparable dataset was provided by Selvaraj *et al.* (2013), who reconstructed F123 haplotypes using HaploSeq, combining Hi-C data with the HapCUT phasing algorithm. The largest chromosome-spanning blocks from GAMIBHEAR and HaploSeq both span over 99.99% of the phasable genome. The largest block from GAMIBHEAR includes >99.9% of observed variants compared to about 95% of observed variants using HaploSeq, a slight improvement due to the large genomic span covered by GAM NuPs. When downsampling the F123 SNV set to human SNV density, HaploSeq and GAMIBHEAR are still able to generate chromosome-spanning, accurate haplotype blocks, however, only 32% of SNVs are phased in the largest block by HaploSeq, while 99.95% of phased SNVs are contained in the largest haplotype block by GAMIBHEAR (Supplementary Note S9).

Although GAMIBHEAR shows high completeness given its input data even at low coverage, the sparsity of the GAM data itself hinders overall completeness. While in the Hi-C data of Selvaraj et al. (2013) 99.6% of variants were covered by at least one read, in the GAM data set only 64.69% of variants are captured. While the sparsity of GAM data does not challenge the generation of accurate 3D chromatin contact maps (Beagrie *et al.*, 2017), advances in the GAM experimental protocol might overcome this drawback in the future to improve phasing results. Additionally, incorporation of statistical phasing could expand the reconstructed haplotypes to uncovered SNVs.

Our proximity scaling model improves the haplotype reconstruction accuracy by taking genomic distances between SNVs into account. The observed decline in phasing information with increasing distance between SNVs is likely due to the formation of highly interacting genomic regions and organisational chromatin structures such as self-interacting TADs (Mb scale) and higher order metaTADs (Razin *et al.*, 2016; Fraser *et al.*, 2015; Ulianov *et al.*, 2016). The MaxST obtained through this proximity-scaled weighted graph discards potential noise and assigns more importance to more likely co-observations of SNVs within neighbouring genomic regions. This runs the theoretical risk of breaking phasing blocks in situations where the only connecting variants were distant in genomic coordinates. In our analysis, no phasing blocks were broken due to proximity scaling of edge weights.

In summary, GAMIBHEAR enables accurate phasing of GAM data with average SERs (~2%) comparable to those obtained with Hi-C (~1.4%) (Selvaraj *et al.*, 2013; Chaisson *et al.*, 2019). While dedicated experimental techniques such as StrandSeq can yield dramatically lower SERs (Chaisson *et al.*, 2019), application of additional experimental techniques to resolve haplotypes more accurately is often not warranted or not feasible due to limited material or costs involved. While GAMIBHEAR is ultimately intended to be used on human data, no GAM dataset of sufficient size is yet available on human samples. In the meantime, the F123 cell line is well-suited to accurately measure phasing performance due to its known haplotype structure before adapting the algorithm to the characteristics of human genomes. Application of our proximity-scaled graph phasing algorithm on F123 GAM data downsampled to human SNV density suggests that the reconstruction of haplotypes is suitable and well applicable for the use in human data as well.

## 5 Conclusion

Understanding the effect of genetic variation on chromatin conformation and gene regulation is a key question in genomics research. Large consortia, such as the 4D Nucleome project (Dekker *et al.*, 2017), are now bundling resources to address open questions in this field and thus allele-specific analyses of chromatin conformation and other sources of genomic variation are moving increasingly into the spotlight (Cavalli *et al.*, 2019). The recently established GAM method (Beagrie *et al.*, 2017) offers a unique opportunity towards high-resolution allele-specific analyses of chromatin contacts in humans, and GAMIBHEAR provides the necessary algorithmic advances towards generating highly accurate, chromosome-spanning haplotypes from GAM data on human samples in the future.

## 6 Author contributions

JM performed bioinformatic analysis on GAM data and implemented the algorithms and R package. JM, BK, SR and RFS designed the algorithms. RK, GL and AK produced the GAM data. AK generated the F123 reference genome. IIA and AK performed bioinformatic analysis and quality control of GAM data. RFS and AP designed and supervised the project.

## 7 Acknowledgements

The authors thank the Helmholtz Association (Germany) for support. AP acknowledges support from the National Institutes of Health Common Fund 4D Nucleome Program grant U54DK107977. IIA was supported by a Long-Term Fellowship from the Federation of European Biochemical Societies (FEBS). SR is supported by DFG Collaborative Research Center SFB 876, subproject C1. JM is supported by DFG Priority Program SPP2202 “Spatial Genome Architecture in Development and Disease”. Computation has been performed on the HPC for Research cluster of the Berlin Institute of Health.

Conflict of Interest: none declared.

## Supplement

### S1 Benchmark Genome

We use the hybrid mouse embryonic stem cell line F123 as a benchmark system for assessing the quality of reconstructed haplotypes from GAM data. The F123 line was derived from the F1 generation of two fully inbred homozygous mouse strains: *Mus musculus castaneus* (CAST) and *Mus musculus domesticus 129S4/SvJae* (J129) (Gribnau *et al.*, 2003).

In order to derive benchmark haplotypes of F123, whole-genome sequencing (WGS) data of CAST and J129 were downloaded from the European Nucleotide Archive (accession number <u>ERP000042</u>) and the Sequence Read Archive (accession number <u>SRX037820</u>), respectively. WGS reads were trimmed using Cutadapt (Martin, 2011) and mapped to the mouse reference genome mm10 using BWA (Li and Durbin, 2009). To determine the haplotypes of the F123 line, SNVs of both parental strains were identified using bcftools (Li, 2011) and SNVs covered by < 5 reads or quality < 30 were excluded.

With the haplotype structure thus known, this cell line serves as the benchmark for all downstream experiments and analyses.

### S2 GAM Dataset, pre-processing and quality control

1281 individual GAM NuPs of the F123 line were generated from the F123 mESC cell line. The F123 SNVs were N-masked in the mm10 reference genome and reads were mapped using Bowtie2 (Langmead and Salzberg, 2012). Duplicate reads were removed using samtools (Li *et al.*, 2009). After mapping, all BAM files and WGS results underwent standard quality control using FastQC (Andrews, 2010) and multiQC (Ewels *et al.*, 2016). Reads were trimmed using BamUtil (Jun *et al.*, 2015) with function trimBam where necessary.

For quality assessment of each sample, the genome was split into fixed windows of size 50kb. For each NuP *i* and each window *j*, the number of reads *r_ij_* and number of nucleotides covered *c_ij_* were determined using bedtools (Quinlan and Hall, 2010). Windows were then classified as *positive* or *negative* based on *r_ij_* and *c_ij_* as follows: From the coverage *c_i_*. of all windows for NuP *i* the empirical nucleotide coverage distribution *P_i_* was computed. From *P_i_*, the minimum coverage percentile *MCP_i_* was chosen such that every window contains three or more reads. The average *MCP* across all NuPs then determined the sample-specific nucleotide coverage thresholds *t_i_* (in bp) for each NuP. Windows *w_ij_* were called positive iff *c_ij_* > *t_i_*, i.e. if the number of nucleotides covered in each window was greater than the sample-specific threshold and negative otherwise. *Positive* windows flanked by *negative* windows on each side were defined as *orphan* windows.

NuPs selected for further analysis had < 60% orphan windows and > 20,000 uniquely mapped reads. 1123 NuPs (89%) passed these quality thresholds (available under 4D Nucleome Consortium data portal accession number 4DNBSTO156AZ, unique 4DN identifiers in Supplementary Data).

Reads were then counted at known heterozygous SNV positions using samtools mpileup (Li *et al.*, 2009). Because of the frequently low coverage from independent (i.e. non-duplicate) reads at most positions (30% of observed SNVs are covered by 2 or fewer reads, 50% by 5 or fewer reads), we counted an allele as present if it was observed in at least one read at the examined position.

### S3 Dataset Statistics

#### Benchmark genome (F123)

The F123 mouse embryonic stem cell line was derived from a hybrid F1 mouse resulting from the cross of the two inbred, homozygous mouse strains CAST (*Mus musculus castaneus*) and J129 (*Mus musculus domesticus J129*). The parental mouse strains are both fully sequenced, their exclusively homozygous genomic variants with respect to the reference mouse genome mm10, which was derived from the mouse strain C57BL/6, are known. The F1 generation resulting from the cross of CAST and J129 is thus heterozygous at all loci for which their parents have different alleles. Their haplotypes are known, making them an ideal model for benchmarking phasing algorithms. Relative to the mouse reference genome mm10, CAST and J129 show 18,892,144 and 4,778,766 germline variants respectively, in concordance with their estimated evolutionary distance from C57BL/6, 371,000 ± 91,000 years (Goios *et al.*, 2007) and approximately 100 years (Simpson *et al.*, 1997), respectively. After exclusion of 2,200,819 overlapping SNV positions and 1,119,044 SNVs located in genomic regions of low mappability, the F123 reference set contains 18,150,228 variants in total, all of which are heterozygous due to inbreeding of the parental strains. Of those, 15,810,835 variants (87.1%) are located on the CAST parental genome, 2,339,393 (12.9%) on J129. This yields an average SNV density of 1 SNV per 132bp, with a median genomic distance of 56bp.

#### Nuclear profiles

We obtained 1281 GAM NuPs of the F123 mESC cell line (4D Nucleome Consortium data portal accession number 4DNBSTO156AZ), out of which 1123 passed quality screening (see Supplementary Note S2).

We extracted on average 305,377 reads from each NuP, covering 0.171% (± 0.167) of the 18,150,228 heterozygous SNVs per nuclear slice (Figure 2A); exemplary data of genomic regions captured in a single NuP is shown in Figure 2B. Out of all F123 SNVs, 11,741,055 (64.69%) were observed at least once across all 1123 NuPs and 7,605,321 SNVs (41.9%) were observed at least twice (Figure 2C). Due to this sparsity and the fact that homologous chromosome pairs occupy distinct chromosomal territories (Khalil *et al.*, 2007), 96.54% of SNV observations showed counts from only one parental allele within one sample. Thus, we removed observed variants with read counts from both parental alleles without substantial loss of information. Since the slicing of nuclei in the GAM experiments is a random process, a balanced observation ratio of alternative and reference alleles of heterozygous SNVs is expected across all NuPs. We thus additionally removed all 550 variants (0.000045%) which significantly deviated from a balanced representation of reference and alternative alleles (p < 0.05 after Benjamini-Hochberg adjustment, binomial test against 0.5).

### S4 Quality measures for reconstructed haplotypes

We here provide details about the employed measures of completeness and accuracy of the reconstructed haplotypes. The measures were chosen to allow comparison between the conceptually different neighbour and graph phasing algorithms and to allow comparison with existing methods. We calculate all SNV-based metrics per chromosome, relative to the number of phasable SNVs *M_c_* on chromosome *c*, i.e. the number of heterozygous SNVs observed at least once in all 1123 NuPs. Analogously, we calculate all metrics based on genomic range (in bp) relative to the phasable genome per chromosome (distance between leftmost SNV and rightmost SNV in bp). We omit chromosome index *c* for brevity in the definitions below and report means and standard deviations of all measures across chromosomes in the Results section of the main text (Table 1). For the number of phasable SNVs and the size of the phasable genome in bp see Supplementary Table 1 below. For a detailed discussion of GAM sparsity see Results and Discussion in the main text.

**Supplementary Table 1:** Description of the phasable SNV set and genome per chromosome. *Full SNV set* describes the phasable SNV set (*M_c_*) and genome as observed in the F123 cell line. *Subsampled* corresponds to the phasable SNV set and genome after employing the downsampling strategy to mimic SNV density in the human genome (Results section 3.3.2).

| Chr | Phasable SNV set (%) |  | Phasable genomic range in bp (%) |  | Chr | Phasable SNV set (%) |  | Phasable genomic range in bp (%) |  |
| --- | --- | --- | --- | --- | --- | --- | --- | --- | --- |
|  | Full SNV set | Sub-sampled | Full SNV set | Sub-sampled |  | Full SNV set | Sub-sampled | Full SNV set | Sub-sampled |
| chr1 | 941707<br>(63.08) | 127839<br>(8.37) | 192365707<br>(98.41) | 192357993<br>(98.41) | chr11 | 640116<br>(68.96) | 86657<br>(9.10) | 118880794<br>(97.38) | 118875736<br>(97.37) |
| chr2 | 782822<br>(66.01) | 106135<br>(8.60) | 178962141<br>(98.27) | 178952453<br>(98.26) | chr12 | 533072<br>(62.68) | 72156<br>(8.33) | 117018721<br>(97.42) | 116999097<br>(97.39) |
| chr3 | 694766<br>(59.03) | 94652<br>(7.83) | 156938923<br>(98.06) | 156927079<br>(98.06) | chr13 | 588006<br>(64.44) | 79840<br>(8.56) | 117318662<br>(97.42) | 117316342<br>(97.42) |
| chr4 | 741959<br>(66.33) | 100655<br>(8.77) | 153307607<br>(97.96) | 153303916<br>(97.95) | chr14 | 553535<br>(63.08) | 74866<br>(8.41) | 121762587<br>(97.49) | 121740416<br>(97.47) |
| chr5 | 738731<br>(64.94) | 100240<br>(8.54) | 148729017<br>(97.96) | 148722202<br>(97.95) | chr15 | 494179<br>(63.75) | 66861<br>(8.43) | 100886142<br>(96.97) | 100866521<br>(96.95) |
| chr6 | 726343<br>(63.99) | 98125<br>(8.51) | 146535902<br>(97.86) | 146533104<br>(97.86) | chr16 | 455935<br>(60.53) | 61556<br>(7.99) | 95024075<br>(96.75) | 94991322<br>(96.72) |
| chr7 | 687475<br>(67.40) | 93734<br>(9.03) | 142338158<br>(97.87) | 142297146<br>(97.84) | chr17 | 431796<br>(65.11) | 58301<br>(8.66) | 91886960<br>(96.74) | 91869956<br>(96.72) |
| chr8 | 688032<br>(67.27) | 93486<br>(8.88) | 126251015<br>(97.59) | 126226500<br>(97.55) | chr18 | 474538<br>(65.43) | 64371<br>(8.65) | 87601865<br>(96.58) | 87592598<br>(96.57) |
| chr9 | 599812<br>(66.89) | 81359<br>(8.81) | 121492381<br>(97.51) | 121483172<br>(97.50) | chr19 | 303591<br>(68.90) | 41486<br>(9.16) | 58235870<br>(94.81) | 58184777<br>(94.71) |
| chr10 | 664640<br>(64.07) | 90115<br>(8.44) | 127492655<br>(97.55) | 127483223<br>(97.54) | **sum** | **11741055**<br>**(64.69)** | **1553527**<br>**(8.56)** | **2403029182**<br>**(97.58)** | **2402723553**<br>**(97.56)** |

#### Completeness and contiguity measures

As a first measure of completeness we report the proportion of heterozygous SNVs and the proportion of neighbouring transitions that have been successfully phased. Because these measures do not take the contiguity of the phased blocks into account we additionally employ metrics that assess the size of the reconstructed haplotype blocks: the S50 (Lo *et al.*, 2011), N50 (Lander *et al.*, 2001) and AN50 (Lo *et al.*, 2011) metrics. A graphical explanation of the completeness measures S50, N50 and AN50 is shown in Supplementary Figure 1A.

##### Number of phased SNVs / transitions

The absolute number of phased SNVs *m_phased_* and its relative counterpart *p_phased_* = *m_phased_*/*M* give a general overview of phasing completeness. In the case where phasing yields a large number *K* of small, disconnected haplotype blocks, the number of phased SNVs will be high, but the phase between these independent blocks is unknown. To account for this fragmentation, we report *t_phased_*, the frequency of phased transitions between adjacent SNVs. As the number of transitions is equal to the number of SNVs *M* – 1 and each additional block beyond the first incurs one unphased transition, this yields:

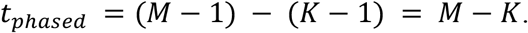

##### S50

S50 (Lo *et al.*, 2011) is a measure of contiguity, i.e. of the size distribution of phased haplotype blocks. To obtain the S50 value, all phased haplotype blocks are sorted by their size (number of SNVs phased in the block), and the S50 value is the size of the block at which 50% or more of SNVs are phased. For example, an S50 value of 1000 SNVs would mean that 50% of all SNVs are contained in haplotype blocks of size 1000 SNVs or larger.

##### N50 / AN50

Analogously, to obtain the N50 contiguity metric (Lander *et al.*, 2001), the phased haplotype blocks are sorted by their genomic span (in bp) to determine the span at which 50% or more of the phasable genome is phased. To correct for cases where isolated haplotype blocks are contained within larger blocks spanning them (see Supplementary Figure 1A), we also report the adjusted N50 (AN50, (Lo *et al.*, 2011)), where the genomic span of the block is adjusted by the fraction of SNVs phased within. Since haplotype blocks reconstructed by the neighbour phasing approach are never nested (Supplementary Figure 1B), N50 = AN50 for neighbour phasing.

##### Accuracy measures

To assess the accuracy of the reconstructed haplotypes we compare GAMIBHEAR estimates with the haplotypes of the F123 mouse embryonic stem cell (mESC) line obtained from whole-genome sequencing of the parental mouse strains (see Supplementary Note S1 ‘Benchmark genome (F123)’). Two measures are considered: the global haplotype agreement calculated by direct comparison of the reconstructed and true haplotypes (i.e. alt-ref configurations) as a global measure of accuracy, and the Switch Error Rate (SER) as a local measure of accuracy (see also Supplementary Figure 1B).

##### Global haplotype accuracy

We report as global haplotype accuracy the overall agreement between haplotypes assigned to SNVs after phasing and their true assignment (see Supplementary Note S1). Let *ĥ* ∈ {−1, 1}^*M*^ be the inferred haplotype assigned to SNV at position *i* and let *h* ∈ {−1, 1}^*M*^ be the true haplotype assignment. The phasing error *e* is then defined as:

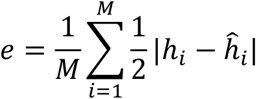

Since the true parent of origin of a variant cannot be identified, haplotypes are equivalent to their full complement in terms of phasing accuracy (for example *h* = (−1, 1, −1) is equivalent to *h′* = (1, −1, 1)). Global accuracy can thus never drop below 50% and the final global haplotype accuracy *G* is thus:

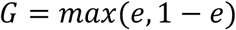

As the first SNV of every haplotype block is arbitrarily set to *h*_1_ = 1, global accuracy for phasing results with many blocks is highly sensitive to the true distribution of alternative alleles over the parental haplotypes. In the F123 dataset, 87 % of alternative alleles reside on the CAST haplotype. A naive phasing algorithm, which places all alternative alleles on haplotype 1 would thus yield a global accuracy of *G* = 0.87. This is visible in the seemingly high *G* = 0.86 of the neighbour phasing algorithm despite its low completeness and contiguity.

##### Switch Error Rate (SER)

We report the Switch Error Rate (SER, Supplementary Figure 1B) as a local accuracy metric. Analogous to the global haplotype accuracy, the SER is defined as the proportion of adjacent variant pairs that were phased incorrectly out of all phased variant pairs. For each haplotype block *k* ∈ {1,…, *K*} we transform the inferred and true haplotype vectors *ĥ*(*k*) ∈ {-1, 1}^*M_k_*^ and *h*(*k*) ∈ {-1, 1}^*M_k_*^ into the inferred vector 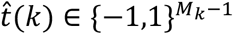 and true transition vector *t*(*k*) ∈ {−1,1}^*M*_*k*−1_^, where 1 and −1 correspond to *stay* and *flip* transitions, respectively. The SER across all haplotype blocks *k* is thus

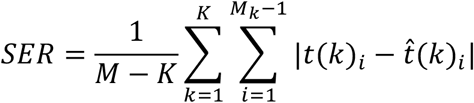

with: 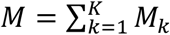 (total number of phased SNVs).

The factor 1/(*M* – *K*) makes the SER relative to all phased transitions.

##### Adjusted SER

Transitions without phasing information that are arbitrarily assigned a *stay* or *flip* state have a 50% chance of being correct, irrespective of the true distribution of alternative alleles over the parental haplotypes. To account for this, we add a SER penalty of 0.5 per unphased transition to define the adjusted SER:

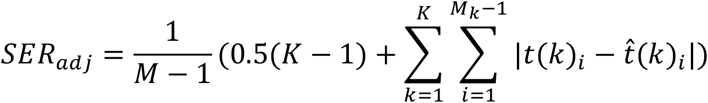

**Supplementary Figure 1:**
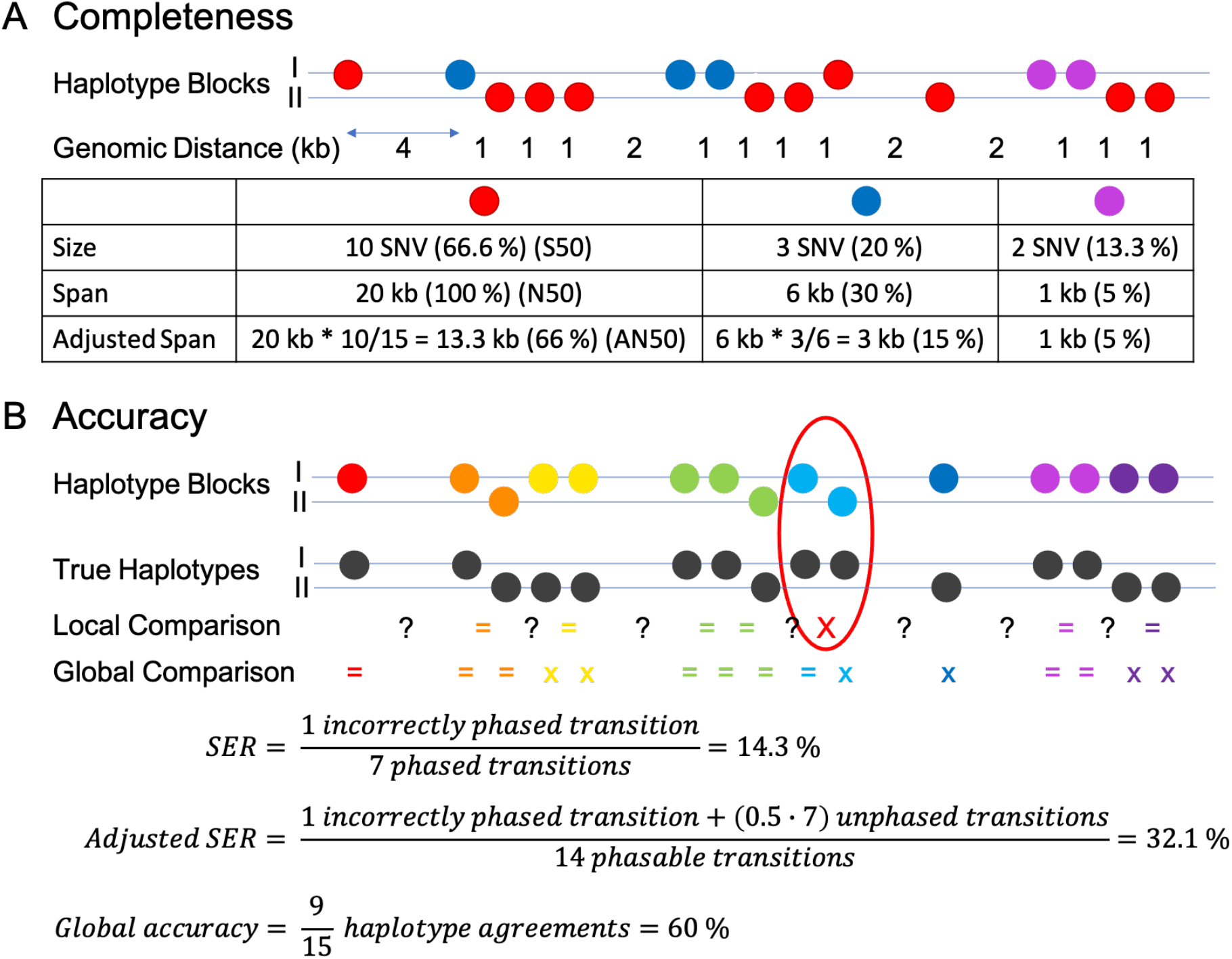
Graphical explanation of quality measures. **A) Completeness:** a schematic graph phasing result is shown, consisting of 3 nested haplotype blocks (red, blue, purple). Size (number of SNVs), genomic span and genomic span adjusted by the fraction of phased SNVs within are exemplarily calculated for the 3 blocks respectively. When ordering the blocks by size, the red block contains more than 50% of the SNV set and thus its size corresponds to the reported S50, N50, AN50 respectively. **B) Accuracy:** a schematic neighbour phasing result is shown, consisting of multiple non-overlapping haplotype blocks. Switch Error Rate (SER) is calculated for the presented haplotype reconstruction, one out of 7 phased transitions is incorrect (circled, marked with a red X). The SER is then adjusted by unphased transitions (marked as ?), which are penalized by 0.5 switch errors. Global Comparison of assignments of alternative alleles to parental haplotypes shows 9 concordant (=) and 6 dissenting (x) assignments, resulting in a global accuracy of 60%.

#### S5 Reconstruction accuracy per chromosome

Here we report global and local accuracy (SER) results of haplotypes reconstructed using the neighbour phasing, basic and proximity scaled graph phasing algorithms per chromosome in Supplementary Table 2. In general, the global accuracy of reconstructed haplotypes improves with the complexity of the used algorithms. Noticeably, the SER metric shows a smaller range in results over chromosomes compared to the global accuracy, which shows outliers. In general, SER is a more meaningful metric compared to global haplotype accuracy, where a single switch error in a haplotype block can lead to the following part of the haplotype block being assigned to the opposite haplotype, thus drastically decreasing global accuracy while maintaining high local accuracy. Supplementary Figure 2 shows one such example of the high impact of switch errors on global haplotype accuracy on an outlier result on chromosome 17.

**Supplementary Table 2:** Global Accuracy (GA) and switch error rate (SER) per chromosome. Haplotypes were reconstructed from the full data set of 1123 GAM NuPs using the neighbour phasing, graph phasing and scaled graph phasing approach. Final haplotype assignments, independent of haplotype blocks, are compared with the known F123 haplotypes. Percent of concordant haplotype assignments (GA, higher is better) and switch error rates of phased transitions (SER, lower is better) are shown per chromosome, as well as mean, median and standard deviation over chromosomes.

| Chr | Neighbour phasing |  | Graph phasing |  | Scaled graph phasing |  |
| --- | --- | --- | --- | --- | --- | --- |
|  | GA | SER | GA | SER | GA | SER |
| chr1 | 85.51 | 0.71 | 95.14 | 5.31 | 98.03 | 1.98 |
| chr2 | 87.22 | 0.74 | 94.92 | 5.80 | 98.00 | 2.08 |
| chr3 | 87.69 | 0.70 | 94.16 | 6.22 | 97.64 | 2.20 |
| chr4 | 84.55 | 0.99 | 94.19 | 6.40 | 97.32 | 2.49 |
| chr5 | 88.29 | 0.74 | 95.44 | 5.12 | 87.44 | 2.03 |
| chr6 | 87.50 | 0.67 | 95.93 | 4.64 | 98.08 | 1.89 |
| chr7 | 85.26 | 0.84 | 95.75 | 5.01 | 97.99 | 2.13 |
| chr8 | 79.99 | 0.93 | 95.44 | 5.28 | 84.51 | 2.29 |
| chr9 | 84.85 | 0.83 | 94.94 | 5.90 | 97.78 | 2.25 |
| chr10 | 93.56 | 0.51 | 95.87 | 4.56 | 98.45 | 1.53 |
| chr11 | 88.90 | 0.68 | 95.20 | 5.49 | 98.14 | 1.87 |
| chr12 | 85.50 | 0.81 | 94.85 | 5.60 | 85.69 | 2.25 |
| chr13 | 87.98 | 0.63 | 95.409 | 5.07 | 98.11 | 1.88 |
| chr14 | 78.16 | 0.92 | 95.505 | 5.12 | 97.65 | 2.33 |
| chr15 | 81.04 | 0.88 | 93.85 | 6.11 | 97.58 | 2.36 |
| chr16 | 88.71 | 0.54 | 95.42 | 5.01 | 98.14 | 1.76 |
| chr17 | 83.718 | 0.93 | 95.16 | 5.60 | 64.79 | 2.54 |
| chr18 | 85.19 | 0.74 | 95.39 | 5.22 | 97.96 | 1.98 |
| chr19 | 87.84 | 0.68 | 94.92 | 5.51 | 97.99 | 1.93 |
| **mean** | **85.87** | **0.76** | **95.13** | **5.42** | **94.28** | **2.09** |
| **median** | **85.51** | **0.74** | **95.20** | **5.31** | **97.96** | **2.08** |
| **sd** | **3.53** | **0.13** | **0.57** | **0.50** | **8.45** | **0.26** |

**Supplementary Figure 2:**
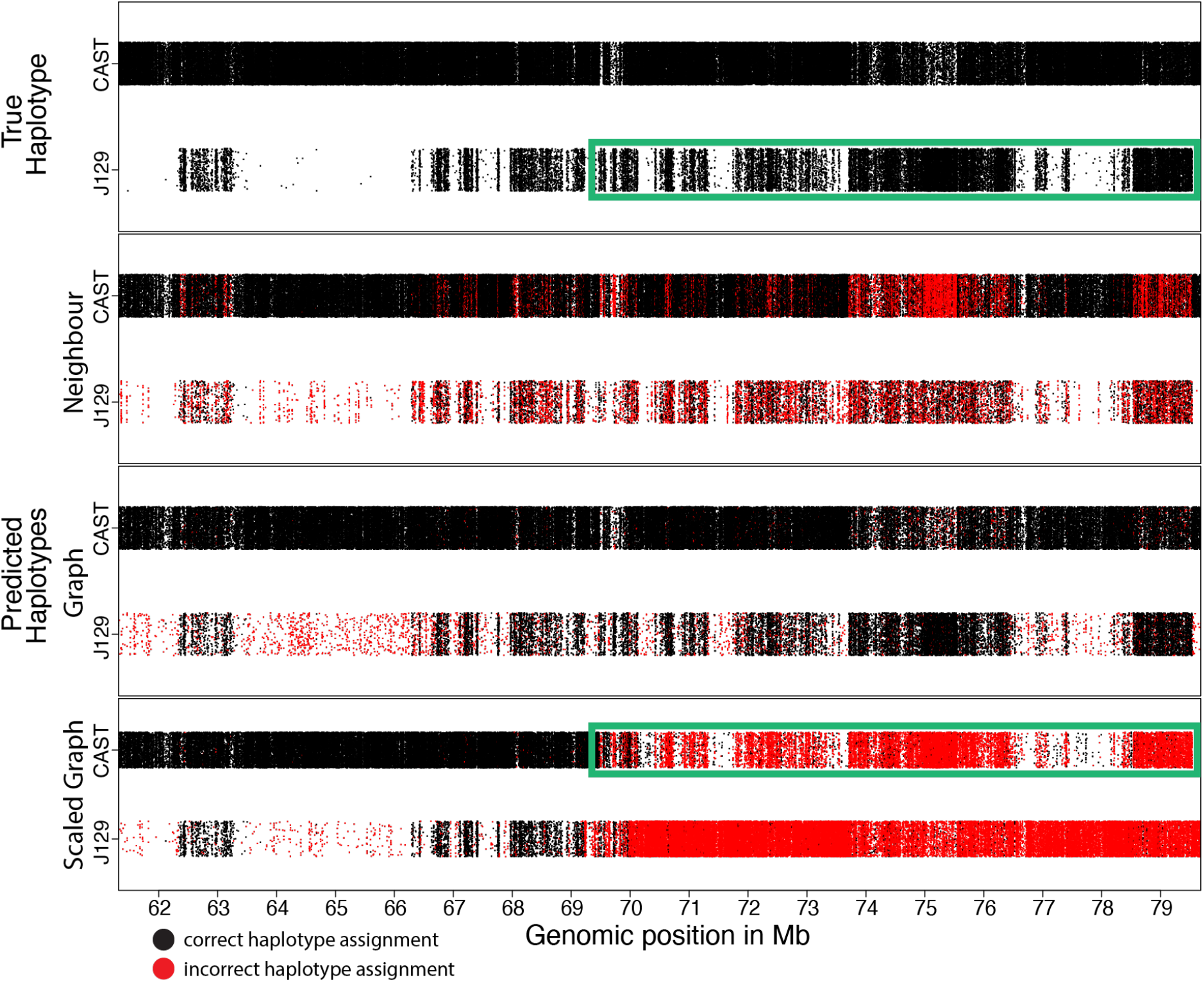
Outlier of decreased global accuracy caused by switch error on chromosome 17. The 4 panels show the location of alternative alleles of heterozygous SNVs along the parental chromosome copies CAST (upper band) and J129 (lower band) in a genomic region on chromosome 17 (61Mb - 80Mb). Each dot represents one SNV, the majority of SNVs is located on the CAST chromosome copies. Panel 1 shows the true haplotypes, meaning the true location of alternative alleles on the parental chromosome copies. Panels 2-4 show the predicted haplotype assignments reconstructed from neighbour phasing (panel 2), graph phasing (panel 3) and scaled graph phasing (panel 4). Black dots represent correct assignments, red dots show incorrect assignments of haplotypes. Graph phasing shows a clear improvement in global accuracy compared to neighbour phasing results. The last panel shows the impact of switch errors within haplotype blocks on the global accuracy: a switch error between 69Mb and 70Mb causes the subsequent haplotype assignment to switch onto the opposite haplotype, causing reduced global accuracy while maintaining local accuracy. The pattern formed by the distribution of J129 SNVs (as shown in panel 1) is still clearly visible in panel 4 after the switch error, only incorrectly predicted to be located on the CAST chromosome copy (highlighted in green boxes), thus demonstrating that the local phasing prediction is still highly accurate.

#### S6 Effect of window size in graph phasing approach

Unless indicated otherwise, all reported results are haplotype reconstructions using default parameter settings. In the graph phasing approach haplotypes are not reconstructed chromosome wide, but in overlapping windows of (default 20,000) SNVs in order to ensure successful completion of calculations without exceeding time and memory limits. We tested the impact of changing window sizes on the quality of reconstructed haplotypes as well as time and memory usage from a minimum of 10,000 SNVs to a maximum of 40,000 SNVs per window. Reducing or increasing the window size only marginally affected the performance of the algorithm in terms of completeness or accuracy; however, it did show a definite impact on the runtime and memory usage (See Supplementary Table 3. Thus, changes to the default parameters should be made with care under consideration of local memory capacity.

**Supplementary Table 3:**
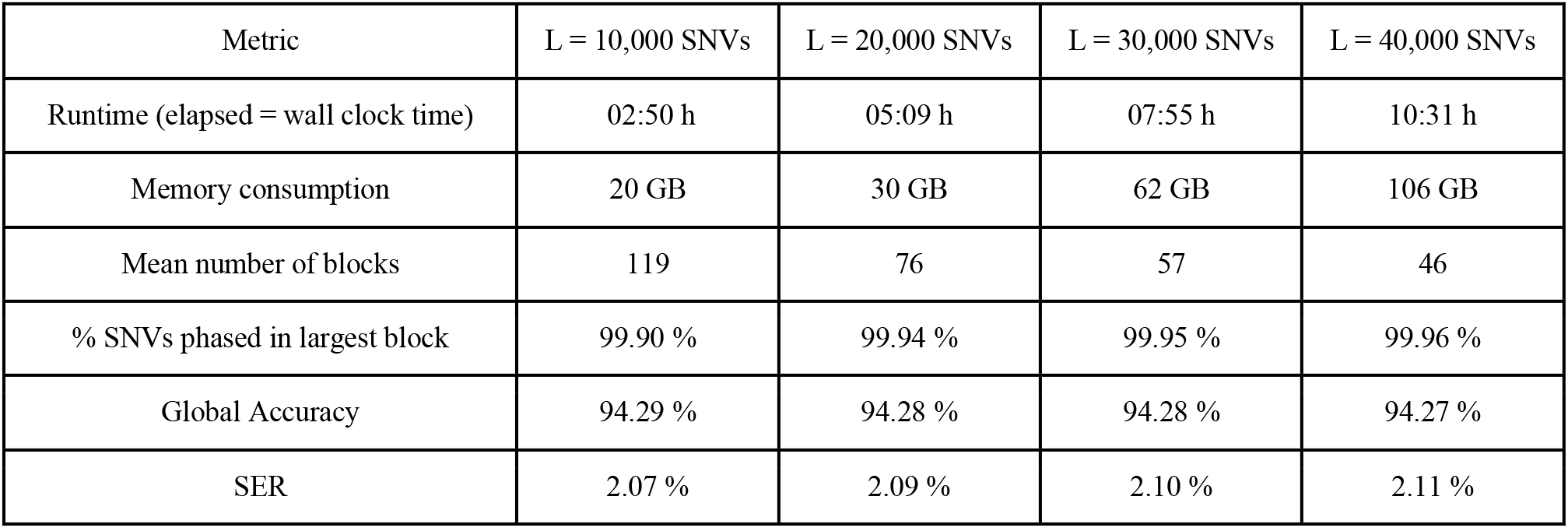
Comparison of different window sizes L. Concerning runtime, memory consumption of the algorithm, as well as completeness and accuracy of reconstructed haplotypes. Scaled graph phasing was used to reconstruct haplotypes from the full dataset.

| Metric | L = 10,000 SNVs | L = 20,000 SNVs | L = 30,000 SNVs | L = 40,000 SNVs |
| --- | --- | --- | --- | --- |
| Runtime (elapsed = wall clock time) | 02:50 h | 05:09 h | 07:55 h | 10:31 h |
| Memory consumption | 20 GB | 30 GB | 62 GB | 106 GB |
| Mean number of blocks | 119 | 76 | 57 | 46 |
| % SNVs phased in largest block | 99.90 % | 99.94 % | 99.95 % | 99.96 % |
| Global Accuracy | 94.29 % | 94.28 % | 94.28 % | 94.27 % |
| SER | 2.07 % | 2.09 % | 2.10 % | 2.11 % |

#### S7 Lower SNV density

The F123 mESC cell line has a relatively high SNV density (8 SNVs per 1kb) compared to humans (approximately 1-1.5 SNVs per 1kb, (1000 Genomes Project Consortium *et al.*, 2015)). To show the effect of SNV density on the quality of haplotype reconstructions, we randomly subsampled the F123 SNV set to resemble human SNV density and evaluated the resulting haplotypes. In order to obtain an average SNV density of 1 SNV per 1kb, we retained 2,462,745 (13.57%) out of the known 18,150,228 F123 SNVs in the 2.46 billion bp mm10 mouse reference genome. The distribution of SNVs along the parental chromosomes remained constant (full SNV set: 87.11% CAST, 12.89% J129; subsampled: 87.14% CAST, 12.86% J129). Variants were randomly subsampled from the true parental haplotypes irrespective of their observation in the GAM NuPs. Similar to the full dataset (64.69% of known SNVs observed), 64.66% of all SNVs were observed in the subsampled dataset. We explored accuracy and completeness of the best-performing proximity-scaled graph phasing algorithm on the subsampled dataset. All parameters, including the proximity scaling parameters, remained unchanged for the haplotype reconstruction. Despite the reduced SNV density and thus increased genomic distance between co-observed SNVs, GAMIBHEAR reconstructed accurate, dense, chromosome-spanning haplotypes: 99.96% of input SNVs were phased into haplotype blocks of minimum size 2, on average 99.95% (± 0.0096%) of those were phased in the main, chromosome-spanning haplotype block, covering 100% (± 0.00%) of the phasable genome.

The mean global accuracy of 87.46% is still fairly high, the high standard deviation of ± 15.21% indicates a large span in the results. The median global accuracy of 96.64% and the switch error rate of 4.84% (± 0.6%) show that the quality of the reconstructed haplotypes in a subsampled dataset is only slightly different from that of the haplotypes reconstructed from the full dataset, indicating that the algorithmic approach is largely independent of SNV density and thus applicable to human data.

#### S8 Comparison with MEC solvers WhatsHap and HapCHAT

GAMIBHEAR is the first algorithm specialized in the usage of GAM data for haplotype reconstruction. GAM data stores phasing information differently than Hi-C data or PacBio long reads, which are frequently used for haplotype reconstruction with existing phasing algorithms such as HapCUT2 (Edge *et al.*, 2017), WhatsHap (Patterson *et al.*, 2015), HapCol (Pirola *et al.*, 2016) and HapCHAT (Beretta *et al.*, 2018).

Chimeric reads from Hi-C experiments store phasing information if both chimeric parts overlap at least one SNV each. If this is the case, phasing information between these two genomic regions is captured, as intrachromosomal contacts are more likely than interchromosomal contacts. On the other hand, reads generated by the PacBio platform capture phasing information regarding all SNVs covered by one individual long read. When it comes to reads generated in GAM experiments, the phasing information is not stored within individual reads as it is the case with chimeric Hi-C reads or long reads, but the SNVs covered by all reads captured within one nuclear profile (NuP) convey accurate local phasing information.

In order to explore if the spatial phasing information from GAM data could be readily transformed for the use in existing phasing algorithms, we decided to transform GAM data into pseudo long reads, since reads in GAM NuPs are sequenced from strands of DNA captured in physical nuclear slices. Thus, all SNVs co-observed within one GAM NuP were treated as if they were all covered on one continuous long read.

For the following comparison we concentrated exclusively on chromosome 1 of the F123 GAM dataset. It contains 1,584,837 known heterozygous SNVs, of which 941,707 are observed in 1110 GAM NuPs. 1087 NuPs contain at least 2 observed SNVs, which is the minimum number of SNVs necessary to convey phasing information. These NuPs were transformed into 1087 pseudo long reads, each read covering between 2 and 25,776 SNVs (on average 2,486 SNVs). Similar to the ternary *N×M* matrix *D* described in section 2.1, *D′* was created from the 1087 pseudo long reads covering 941,704 SNVs in total. *D′* was built to meet the tools’ internal representation of reference, alternative and not observed alleles as {0, 1, −}, respectively, and then used as direct input to the wMEC and k-constrained MEC solvers WhatsHap and HapCHAT, using default parameters.

WhatsHap is fixed parameter tractable in the coverage and sets a default coverage threshold of 15x (maximum 23x) since PacBio long read data is characterized by uniform read and SNV coverage. However, unlike in PacBio long read data, SNV coverage in the sparse GAM data is usually low (chr1: on average 2.87x) but not uniform and varies, up to 37x on chr 1.

Thus, a few SNVs in *D′* (0.23%) exceed the default coverage of 15x, and 0.0073% exceed even the maximum coverage threshold (23x).

To ensure the compliance of its coverage threshold, WhatsHap uses a read selection heuristic to select suitable reads that are most informative for phasing. The read selection process of WhatsHap resulted in a loss of the majority of long reads, 69 reads (6.35%) remained. As the coverage of GAM data is usually low and most SNVs are only observed once, this stringent read selection resulted in a loss of the majority of considered SNVs. 11,039 SNVs (1.17% of input SNVs) were retained for subsequent phasing, but haplotypes containing approximately 1% of input SNVs would not be useful. In conclusion, as WhatsHap’s read selection heuristic was designed for data sets where SNV coverage is uniform, GAM data cannot be readily transformed for its use in WhatsHap as it does not meet coverage requirements.

HapCHAT, based on WhatsHap and HapCol, was precisely developed to allow the consideration of datasets composed of higher coverages, as well as to improve the accuracy of computed haplotypes. In a preprocessing step, reads that are likely to originate from the same chromosome copy are merged. It was shown that, using read merging, HapCHAT can effectively handle datasets with approximately 60x coverage.

In our comparison, the 1087 pseudo long reads were merged into 691 reads. To fulfill the default coverage threshold of 15x, merged reads were downsampled using the same selection process as employed by WhatsHap. The 63 remaining merged pseudo long reads covered 604,358 SNVs (64.18% of input SNVs), all of which were subsequently phased into 5 haplotype blocks of minimum size 2. The largest block contained 604,350 SNVs (S50, 64.18%) and spanned 192,334,685 bp (99.993% of the phasable genomic range), creating a chromosome-spanning haplotype block. Adjusting its span for the fraction of phased SNVs yields 123,434,304 bp (AN50), which is equivalent to 64.17% of the phasable genomic range.

The haplotypes reconstructed by HapCHAT, which eventually phased 64.18 % of input SNVs show a global accuracy of 81.36%, with a SER of 11.38%. The MEC cost was reported as 307,734. A side-by-side comparison is shown in Supplementary Table 4.

**Supplementary Table 4:** **Side-by-side comparison of HapCHAT and GAMIBHEAR** phasing results on F123 chromosome 1.

| Metric | HapCHAT |  | GAMIBHEAR |  |
| --- | --- | --- | --- | --- |
|  | Absolute | Percent | Absolute | Percent |
| Phased variants | 604,358 SNVs | 64.18 % | 941,400 SNVs | 99.97 % |
| Number of Blocks | 5 | - | 125 | - |
| S50 | 604,350 SNVs | 64.18 % | 941,060 SNVs | 99.93 % |
| N50 | 192,334,685 bp | 99.99 % | 192,348,818 bp | 100 % |
| AN50 | 123,434,304 bp | 64.17 % | 192,216,665 bp | 99.93 % |
| Global Accuracy | - | 81.36 % | - | 98.03 % |
| SER | - | 11.38 % | - | 1.98 % |

#### S9 Comparison with HaploSeq

In 2013 Selvaraj et al. presented HaploSeq, a method that combines Hi-C on the experimental side, and HapCUT on the algorithmic side, to reconstruct accurate haplotypes genome-wide. HaploSeq, as well as GAMIBHEAR, were developed and validated on the F123 mESC line, which makes their results comparable. The authors of HaploSeq present quality metrics with respect to the largest haplotype block, defined as the block with the most variants phased (MVP block). To enable direct comparison between the approaches, we report here the same metrics as reported by Selvaraj et al (see Supplementary Table 5). Since GAMIBHEAR was developed to enable haplotype-specific analysis of GAM data without the need of further experiments, we primarily report our results with respect to the set of SNVs observed in the GAM data set, but additionally with respect to the full set of known heterozygous SNVs in F123.

Selvaraj et al., 2013 report > 99.9% of each phasable chromosome spanned by the MVP block, phasing about 95% of variants into the largest block, and > 99.5% accurately phased SNVs.

**Supplementary Table 5:**
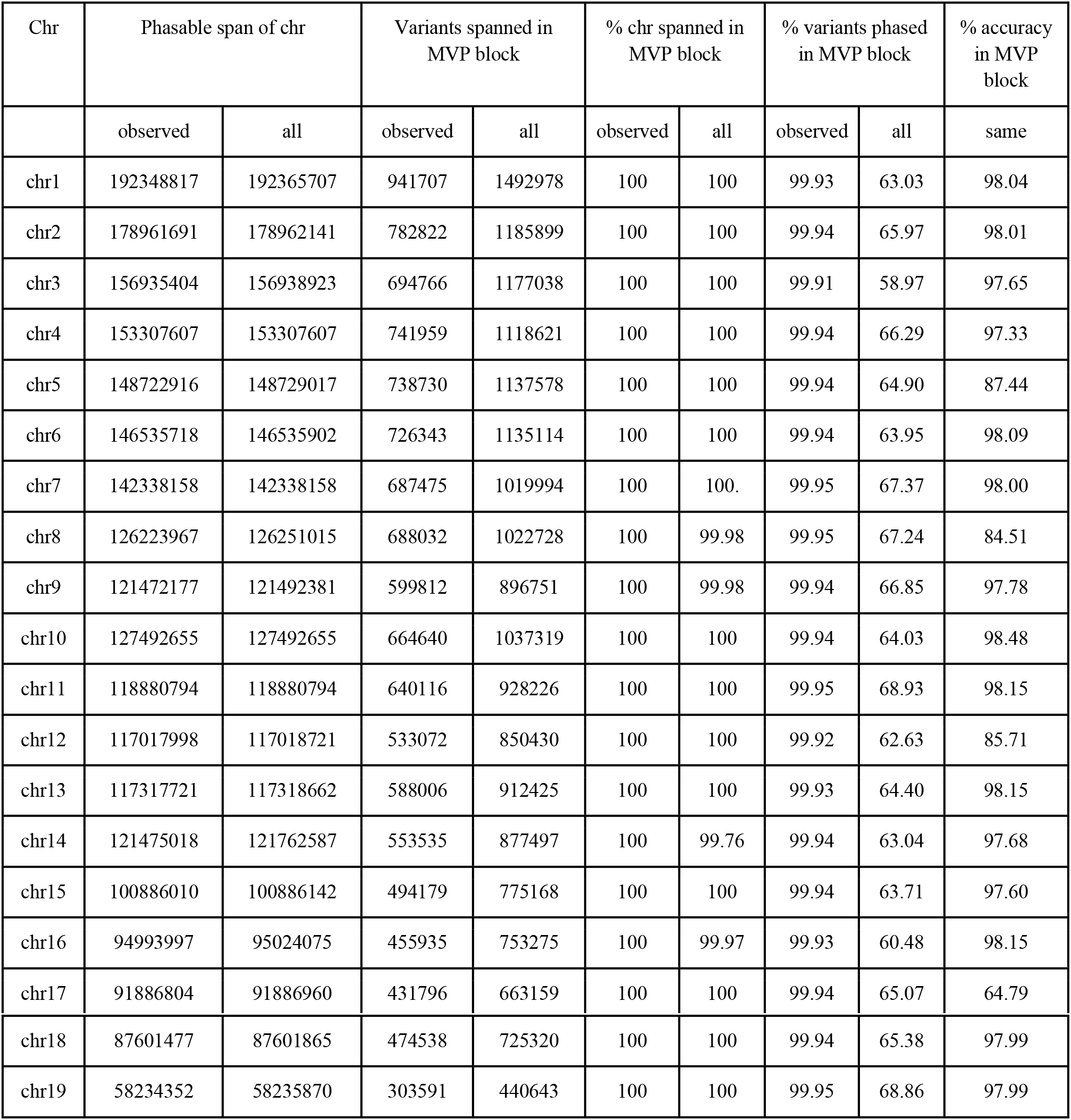
GAMIBHEAR metrics comparable to HaploSeq.

When downsampling the F123 SNV set to human SNV density, Selvaraj et al. still report complete (>99.2 % of the phasable chromosomes spanned) and only marginally less accurate (> 98.9%) MVPs, which, however, show a drastically lower resolution as the MVP blocks only phased approximately 32 % of SNVs.

In contrast, GAMIBHEAR was able to phase 99.96% of downsampled input SNVs, of which 99.95% are contained within the main, chromosome-spanning haplotype block. This block spans 100% of the phasable genome (97.56 % of the full genome) with a comparable global accuracy of 96.64%.

